# Epithelium Stratifies via Nucleation and Growth Induced by Foam-Geometric Instability

**DOI:** 10.1101/2024.12.26.630435

**Authors:** Shuya Fukamachi, Razib Datta, Duligengaowa Wuergezhen, Takehiko Ichikawa, Rei Yagasaki, Phoebe Leeaw, Makiko Arai, Aki Teranishi, Shuhei A. Horiguchi, Kohei Omachi, Aiko Sada, Itaru Imayoshi, Kentaro Kajiwara, Tetsuya Hiraiwa, Masanobu Oshima, Takeshi Fukuma, Hironobu Fujiwara, Satoru Okuda

## Abstract

The epithelium undergoes stratification, transitioning from a monolayer to a multilayer structure, across broad phenomena. Recent studies have identified several cell behaviors as triggers, including junctional tension, cell density and geometry, and topological defects. However, how these factors drive stratification throughout the entire epithelium remains poorly understood. Here, we report a mechanism underlying epithelial stratification that mirrors the physics of phase transition. Combining cell culture with three-dimensional vertex modeling, we demonstrate that epithelial stratification is analogous to a structural phase transition driven by the nucleation-growth process, i.e., multilayer origins dispersedly appear and expand across the epithelium via unordered intermediate states. This transition is induced by a mechanical instability inherent in the foam-like geometry of the epithelium. Moreover, the nucleation-growth concept applies to embryonic skin development and intestinal cancer transformation. These findings conceptualize epithelial stratification as a form of a phase transition governed by foam mechanics, offering a physical perspective on various epithelial developments.

## Introduction

Many epithelia develop multilayer sheet structures^1^, such as those found in the skin^2^, hair follicles^3^, cornea^4^, esophagus^5^, stomach^6^, intestine^7^, bronchus^8^, and brain^9^. These multilayer structures typically evolve from a monolayer through a stratification process, a key phenomenon in embryogenesis. Defects in this process lead to fatal diseases. For instance, during embryogenesis, mouse skin, which normally forms a multilayer, remains a monolayer when the p63 gene is defective, resulting in postnatal death due to dehydration^10–12^. Moreover, during carcinogenesis, abnormal cells undergo stratification to form tumors on epithelial sheets^13^. Consequently, epithelial stratification has been widely studied in various biological contexts. However, although key molecules and cell behaviors have been identified, the mechanism of collective cell dynamics inducing the stratification of the entire epithelium remains poorly understood. Thus, it is crucial to develop a comprehensive understanding of the stratification process, particularly how individual cell delaminations lead to the stratification of the overall epithelium.

During epithelial stratification, cells typically delaminate from either the apical or basal side and accumulate vertically within the epithelial sheet. Several mechanisms have been proposed to explain delamination of individual cells. For example, in skin and brain development, cell delamination often results directly from vertically oriented cell divisions^14–16^, whereas recent pioneering work has shown that cells stratify even without the regulation of cell division orientation^15^. Additionally, cells that lose their basal side during mitosis can regain it by elongating their protrusions^17,18^. These findings suggest that transient vertical arrangements by oriented cell divisions may not be sufficient; instead, a balance of mechanical forces among cells is required to form a stable, stratified structure.

From a physical viewpoint, apical junctional forces, such as increased actomyosin contractility and reduced cadherin-mediated adhesion, are well-known contributors to cell delamination^19–24^. Moreover, an increase in cell density within the epithelial plane can drive cells out of the layer^25–27^. Recent studies have also revealed the impact of cell geometry on this process^28–31^; particularly, topological defects trigger initiation of the stratification process^32,33^. These findings underscore the critical role of mechanical forces in epithelial stratification. However, current understanding is largely limited to the initiation of the stratification process, i.e., single-cell delamination, and little is known about how the delamination of individual cells expands into overall tissue stratification. Specifically, it remains unknown whether this transition propagates as a wave originating from a specific point of cell delamination or instead emerges from numerous origins, spreading through an unordered intermediate state, and what physics underlies this transition.

To bridge the gap between cell- and tissue-scale mechanics, we have combined three-dimensional (3D) quantifications and multicellular dynamics modeling. This approach elucidates how individual cells mechanically control the overall layer structure of the epithelium. Two primary objectives guided our study. First, we aimed to observe the collective cell dynamics in 3D space during epithelial stratification. Although the kinetic behaviors of individual cell delamination are well documented, the collective motions of cells throughout the entire stratification process remain largely unexplored. Specifically, the spatial distribution of delaminated cells during stratification and their subsequent influence on the delamination of other cells requires further clarification. Second, we focused on understanding the physical mechanisms underlying collective cell motion during stratification. Given the challenges of monitoring mechanical interactions in 3D tissue structures, we developed a 3D vertex model^34,35^. This model captures the dynamics of stratification, including 3D deformation and cellular arrangements. This approach allows for a comprehensive analysis of the 3D structures and mechanics involved in multicellular arrangements during epithelial stratification.

To elucidate the fundamental mechanisms of epithelial stratification, we observed the stratification process of Madin-Darby canine kidney (MDCK) cells cultured in a dish and employed a 3D vertex model to understand the physical mechanisms of stratification. Although MDCK cell cultures and vertex models have often been used to study cell delamination^36–38^, little has been done to extensively apply them to stratification. In both systems, we recapitulated the kinetic process of epithelial stratification at single-cell resolution, i.e., how individual cells delaminate and accumulate to form multilayer structures. To analyze this process, we quantified 3D cell configurations and their dynamics to reveal the physical concept of the phase transition with nucleation and growth, which could be applied to epithelial stratification. To understand the mechanisms of delamination, we performed analytical calculations using a 3D vertex model to reveal a mechanical instability inherent in the foam-like epithelial geometry. To validate the model predictions, we further conducted image analyses and atomic force microscopy (AFM) measurements. Finally, we explored the applicability of the nucleation-growth concept to embryogenesis and carcinogenesis by observing skin development in mouse embryos and malignant transformation of intestinal cancer organoids. Throughout these interdisciplinary investigations, we conceptualized epithelial stratification based on the physics of phase transition.

## Results

### Epithelial stratification is akin to a phase transition via nucleation and growth

To understand the kinetic process of epithelial stratification, we employed simple MDCK cell cultures on a dish coated with collagen gel (Fig. 1a, see Materials and Methods). Although MDCK monolayers are generally stable, minor multilayering can occasionally occur even under standard culture. In addition, prior studies have shown that non-transformed MDCK cells can form multilayers under specific conditions^39,40^, and that apical stacking on an existing MDCK sheet is mechanically realizable^41^. Under our culture conditions (see Materials and Methods), this propensity was noticeably increased. We therefore used this setting in which the cells spread and stratified on the gel.

**Fig. 1:**
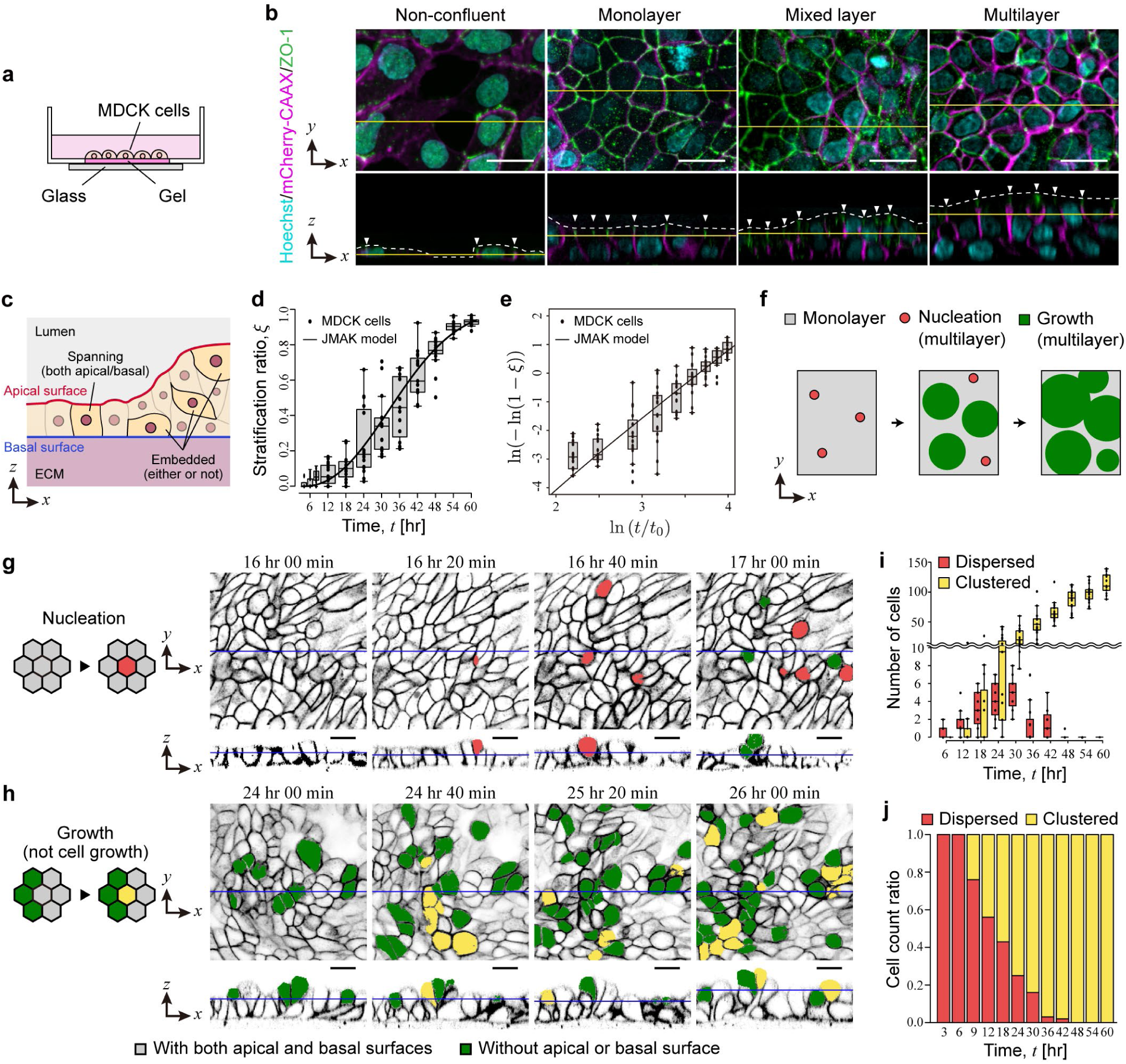
Epithelial cells stratify through a nucleation-growth process. **a** Schematic illustration of the MDCK cell culture setup used to observe epithelial stratification. **b** Fluorescence images of MDCK cells stably expressing mCherry-CAAX, stained with a ZO-1 antibody and Hoechst. The top panel shows the *z*-projection, and the bottom panel displays a single *xz*-view image along the yellow line in the top panel. The white arrow highlights the ZO-1 signal in the *xz*-view, and the dotted line marks the apical surface. Scale bar: 20 µm. **c** Cell categories. Spanning indicates cells have apical and basal surfaces, and embedded indicates cells that lack either the apical surface, basal surface, or both. **d**, **e** Linear and Avrami plots of the stratification ratio, *ξ* (*N* ≥ 5). In (**d**, **e**), solid lines represent fitted curves based on the JMAK model (detailed in Materials and Methods). **f** Schematic illustration of the nucleation and growth process that describes the transition from a monolayer to a multilayer structure. **g**, **h** Schematic illustration (left) and corresponding experimental images (right) depicting nucleation and growth during stratification of MDCK cells. Red cells represent newly delaminated cells appearing in the monolayer, and yellow cells represent newly delaminated cells adjacent to existing ones. Both were classified as lacking apical or basal surfaces. Scale bar: 20 µm. **i** Temporal changes in the number of dispersed versus clustered delaminated cells. **j** Temporal changes in the ratio of dispersed and clustered delaminated cells. Plots in (**d**, **e**, **i**, **j**) include data from 15 images. Box plots in (**d**, **e**, **i**): center: median; bounds, 25th and 75th percentiles; whiskers extend 1.5 times the interquartile range from the 25th and 75th percentiles; outliers are represented by dots.

Under this condition, we identified four distinct stages during this stratification (Fig. 1b): a nonconfluent structure where cells partially occupied the dish, a complete monolayer with cells densely packed on the dish, a mixed layer featuring partially stratified cells, and a multilayer with cells mostly stratified. Throughout the stratification, the tight junction marker ZO-1 consistently localized at the cell-cell boundaries around the apical surface of the entire sheet (Fig. 1b). In contrast, cells delaminating from the apical side lost their junctions. Thus, the cells stratified from a monolayer to a multilayer structure while maintaining the apicobasal polarity of the entire tissue.

To understand this stratification process, we quantified its time evolution (detailed in Materials and Methods). The structure of the epithelium consists of two types of cells: cells that span from the apical to basal surfaces of the epithelium and cells embedded within the epithelium that lack either an apical surface, a basal surface, or both (Fig. 1c). Here, the apical surface refers to the surface of the cell facing the lumen or free space on the apical side of the epithelium, while the basal surface refers to the surface in contact with the extracellular matrix (ECM). Loss of either the apical surface, basal surface, or both leads to delamination.

Based on this classification, we introduced the stratification ratio, *ξ*, defined as the occupancy ratio of the stratified domain on the plane of the dish:

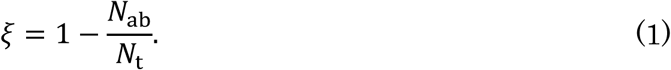

In Equation 1, *N*_ab_ represents the number of cells that span both the apical and basal surfaces of the epithelium, and *N*_t_ represents the total number of cells within the epithelium. We observe a gradual increase in *ξ* from 0 to approximately 1, with an S-shaped curve (Fig. 1d, e), signifying a structural crossover from a complete monolayer to a fully multilayer arrangement.

Next, to understand the kinetic process of individual cell behaviors, we examined their delamination processes. During stratification, individual cells gradually delaminate by losing their apical or basal side, dividing the entire epithelium into two domains: monolayer and multilayer. We identified two characteristic events in stratification: nucleation and growth (Fig. 1f). Nucleation refers to cell delamination events occurring individually in a monolayer domain (Fig. 1g). Growth refers to cell delamination events occurring at the boundary of the monolayer adjacent to a multilayer domain (Fig. 1h). These events mirror the physical processes of nucleation and growth often observed in structural phase transitions, such as crystallization.

To understand the time evolution of the nucleation and growth events in epithelial stratification, we quantified their frequencies. Because it was difficult to directly measure each event, we counted the number of delaminated cells, categorizing them as either dispersed or clustered, i.e., isolated or adjacent to other delaminated cells (detailed in Materials and Methods). Dispersed delaminations initially increased during the early stage of stratification, followed by an increase in clustered cells (Fig. 1i). Consistently, the most frequent size of delaminated cell clusters shifted from single cells to multiple cells (Fig. 1j). Notably, the increase in clustered cells eventually surpassed that of dispersed delaminations. This pattern is reminiscent of the nucleation-growth process of structural phase transitions, characterized by initial frequent nucleation and subsequent growth. This similarity draws an analogy between the mechanisms of epithelial stratification and the phase transition with nucleation-growth process.

Based on the analogy between epithelial stratification and nucleation-growth physics, we applied methods used in the physical understanding of nucleation and growth to investigate the epithelial stratification process. Specifically, we applied the well-known Johnson-Mehl-Avrami-Kolmogorov (JMAK) model^42^ and fitted the time evolution of *ξ*. Originating from a classical nucleation-growth theory, the JMAK model describes the progression of nucleation and growth and is expressed as a simple sigmoid function representing the time variation of the phase proportion:

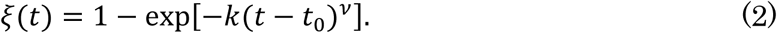

In Equation 2, *k* represents the phase growth rate, *t*_0_ represents the initiation time, and *ν* represents the Avrami exponent. Remarkably, *ξ* values obtained from the MDCK cell culture were successfully fitted using the JMAK model (Fig. 1d, e; detailed in Materials and Methods). The Avrami exponent was determined as *ν* = 2.29. Generally, *ν* reflects the characteristics of the phase transition, and it is equal to 3 for two-dimensional (2D) growth with random nucleation sites. However, *ν* is decreased if the nucleation site distribution is not random, indicating restricted growth. Our obtained *ν* value is reasonable, and lower than 3, suggesting that nucleation is influenced by the already stratified domain. These results demonstrate that epithelial stratification can be conceptualized as a form of a phase transition with nucleation-growth process.

### 3D foam mechanics-based modeling reproduces epithelial stratification

To investigate nucleation-and-growth mechanisms underlying epithelial stratification, we developed a 3D vertex model (detailed in Materials and Methods) based on the assumption that epithelial stratification can be regarded as a mechanical process of foam-like materials. This model overcomes the limitations of conventional models; while 3D particle models and 2D vertex models have unveiled the mechanics of stratification^13,43–45^, they cannot fully describe 3D cell deformations and arrangements. Our model describes a 3D foam geometry of the epithelium equilibrated under stress-free boundary conditions, assuming balanced mechanical forces among cells (Fig. 2a; detailed in Materials and Methods). Cells have apicobasal polarity (Fig. 2b); based on observations of the tight junction marker ZO-1 (Fig. 1b), we postulated that all cell-cell boundary surfaces comprised parts of the basolateral surfaces, and the other cell surfaces composing the tissue surface were defined as the apical surfaces. Moreover, we introduced two types of junctional contractility, well-established regulators of cell delamination: apical perimeter elasticity of each cell, *Γ*, and line tension at apical junctions, *λ* (Fig. 2c). These parameters are known to describe apical behaviors of epithelial cells in 2D space^46^ and have been applied to those in 3D space^47–49^. While *Γ* homogenizes the size of apical cell surfaces across adjacent cells, *λ* works oppositely^30^.

**Fig. 2:**
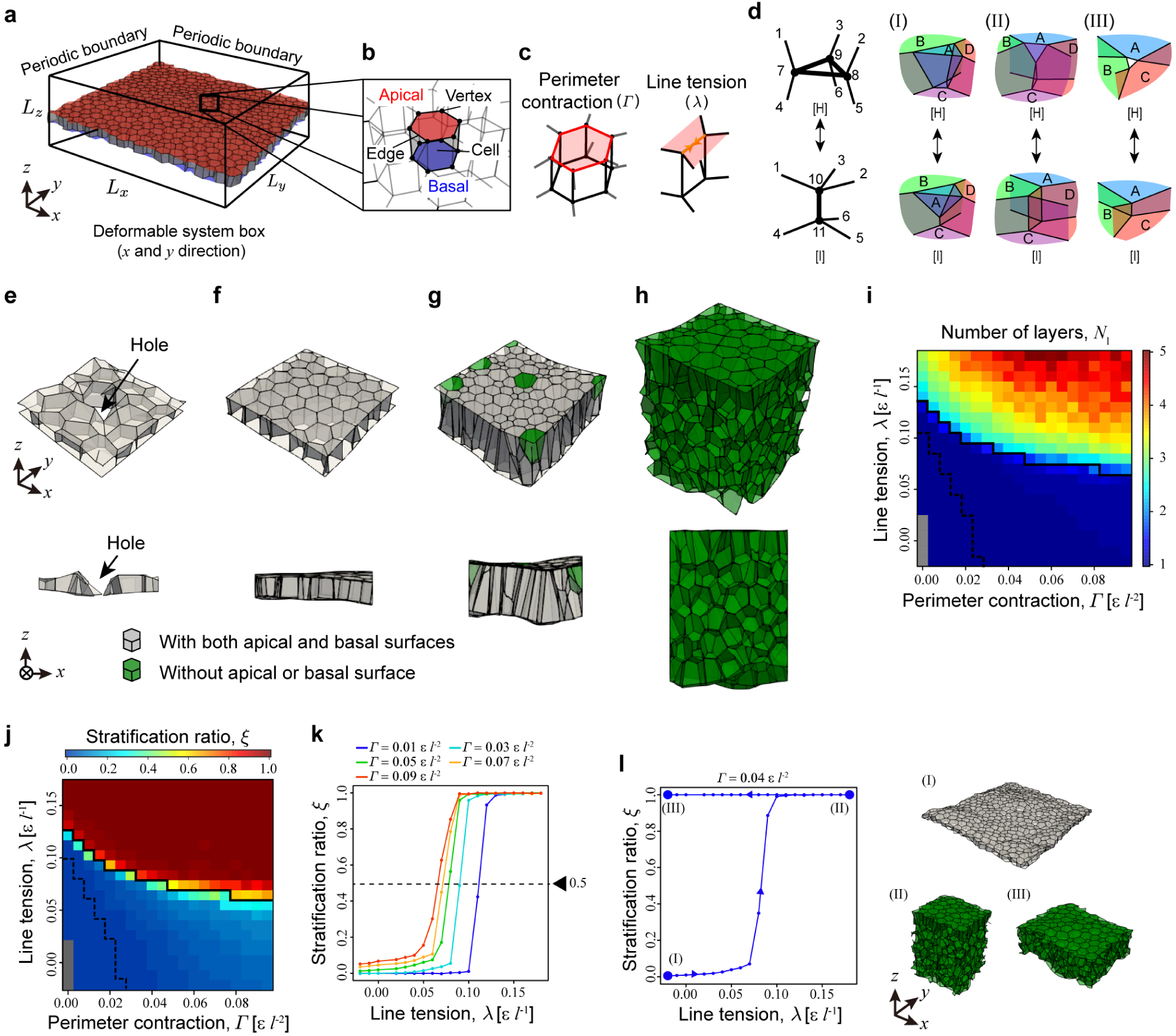
Three-dimensional vertex model reveals a phase transition from monolayer to multilayer phase. **a** Initial monolayer epithelial sheet condition in 3D space, where the system box is deformed by axial stress in the *xy* plane. **b** Enlarged image of a single cell within the epithelial sheet; surface colors indicate the apical (red), lateral (gray), and basal (blue) surfaces. **c** Schematic illustrations of apical junctional forces acting on individual apical cell perimeters: apical perimeter contraction, *Γ* (left), and individual apical cell edges: apical line tension, *λ* (right). **d** Definition of topological operations between patterns [H] and [I], including examples of cell delaminating from a monolayer (I), cell rearrangement within the sheet plane (II), and sheet collapse through hole formation (III). (**e**–**h**) Epithelial structures after relaxation, showing different states: collapse (**e**), monolayer (**f**), partial multilayer (**g**), and multilayer (**h**), induced by varying apical junctional forces. Green cells represent those lacking apical, basal, or both surfaces. The parameters used were set to *Γ* = 0*, λ* = 0 (**e**), *Γ* = 0.015 *ε l*^-2^*, λ* = 0 (**f**), *Γ =* 0.075 *ε l*^-2^*, λ* = 0.03 *ε l*^-1^ (**g**), and *Γ* = 0.065 *ε l*^-2^*, λ* = 0.13 *ε l*^-1^ (**h**). **i** Number of layers, *N*_l_, with respect to *Γ* and *λ*. **j** Stratification ratio, *ξ*, with respect to *Γ* and *λ*. **k** Stratification ratio, *ξ*, as a function of *λ*. **l** Hysteresis behavior in epithelial stratification. Epithelial structures at the indicated points (I)–(III) are shown on the right. The results between points (I) and (II) were calculated starting from the monolayer initial condition shown in (**a**), and those between points (II) and (III) were obtained using the multilayer initial condition at point (II). The embedded images indicate epithelial structures at points (I)–(III). In (**i**, **j**), solid and dotted lines indicate boundaries among the multilayer (*ξ >* 0.5), monolayer (0.5 ≥ *ξ* ≥ 0.1), and complete monolayer (0.1 > *ξ*) phases. Gray-colored regions in (**i**, **j**) indicate collapse states. Colors in (**i**, **j**) indicate the average values of the five initial conditions. The gray-colored regions indicate the collapsed state. Each data point in (**i**–**l**) indicates the average values calculated from the five initial conditions.

The mechanical behavior of cells with apicobasal polarity is given by the effective energy *U*, expressed as

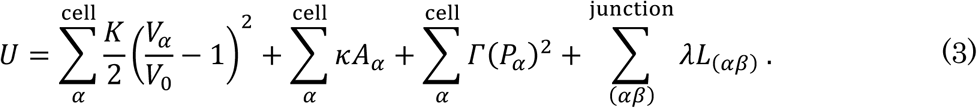

In Equation 3, *K*, *V*_0_, *κ*, *Γ*, and *λ* are constants, and *K*, *V*_0_ *κ*, and *Γ* are positive. *α* and *β* denote cell indices, and (*αβ*) indicates a pair of the *α*th and *β*th cells. The first term represents the volume elastic energy, which restores the volume of each cell *V*_*α*_ to *V*_0_ (set to *l*^3^, where *l* represents the unit length), with the elastic constant *K*. The second term is the basolateral surface energy, characterized by the surface tension, *κ*, which acts to contract the basolateral surface area *A*_*α*_. The third term denotes the elastic energy associated with the apical junctional perimeter, where the elastic constant *Γ* drives contraction of the apical perimeter *P*_*α*_. The fourth term accounts for the tensile energy, where the tension constant *λ* contracts the length of the apical junction between neighboring cells, *L*_(*α*_*_β_*_)_.

Using this model, we numerically calculated the energy relaxation process starting from an initial condition of a complete monolayer (Fig. 2a; detailed in Materials and Methods) and obtained an equilibrium structure at the local energy minimum. During this relaxation, cell configurations were rearranged in 3D space, leading to stratification. This cell rearrangement is described by introducing operations named [H]-[I] (Fig. 2d), as suggested in a previous study.^34^ These operations cause the delamination of a cell from a monolayer to either the apical or basal side (Fig. 2d-I), cell rearrangement within the plane of an epithelial sheet (Fig. 2d-II), and the collapse of the epithelial sheet, creating a hole (Fig. 2d-III).

By exploring the apical junctional parameters *Γ* and *λ*, we replicated all four epithelial structure types: nonconfluent structure (Fig. 2e), complete monolayer (Fig. 2f), mixed layer (Fig. 2g), and multilayer (Fig. 2h), corresponding to those observed in MDCK cell culture (Fig. 1b). We calculated the average number of layers in individual epithelia, which increased with *Γ* and *λ* (Fig. 2i).

To analyze the dependence of the layer structure on the apical junctional parameters *Γ* and *λ*, we calculated the stratification ratio, *ξ* (Fig. 2j). At extremely small values of *Γ* and *λ*, the epithelium formed an incomplete monolayer, which transitioned to a complete monolayer as *Γ* and *λ* slightly increased. As *Γ* increased further, the epithelial structure transitioned from a complete monolayer to a mixed monolayer, and as *λ* increased further, it transitioned to a complete multilayer structure. Quantitatively, *ξ* jumped from 0 to 1 with an increase in *λ* (Fig. 2k), indicating a phase transition from monolayer to multilayer.

Moreover, to investigate the phase transition characteristics, we varied *Γ* and *λ* over time and observed the evolution of *ξ* starting from the initial condition of either a monolayer or a multilayer structure (Fig. 2l). The monolayer-to-multilayer transition was observed when starting from a monolayer (Fig. 2l, from points I to II), but the multilayer-to-monolayer transition did not occur when starting from a multilayer (Fig. 2l, from points II to III). These results show that this transition exhibits hysteresis. Thus, our 3D foam-based model replicates the monolayer-to-multilayer transition observed in epithelial stratification, characterizing it as a first-order-like phase transition.

### The nucleation-growth process drives epithelial stratification

To elucidate the physical process of epithelial stratification, we observed the dynamics of each cell in the numerical simulations shown in Fig. 2. We identified both nucleation (Fig. 3a) and growth (Fig. 3b) events, as observed in MDCK cell culture (Fig. 1g, h). Moreover, to understand the nucleation-and-growth characteristics of the simulated stratifications, we applied the JMAK model to fit the time evolution of *ξ*. Equation 2 is successfully fitted to the *ξ* values obtained from the simulations (Fig. 3c, d). From this fitting, we obtained *ν* values ranging from 1.0 to 2.69 (Fig. 3e). The decrease in *ν* with an increase in *Γ* indicated that nucleation sites became more restricted as *Γ* increased.

**Fig. 3:**
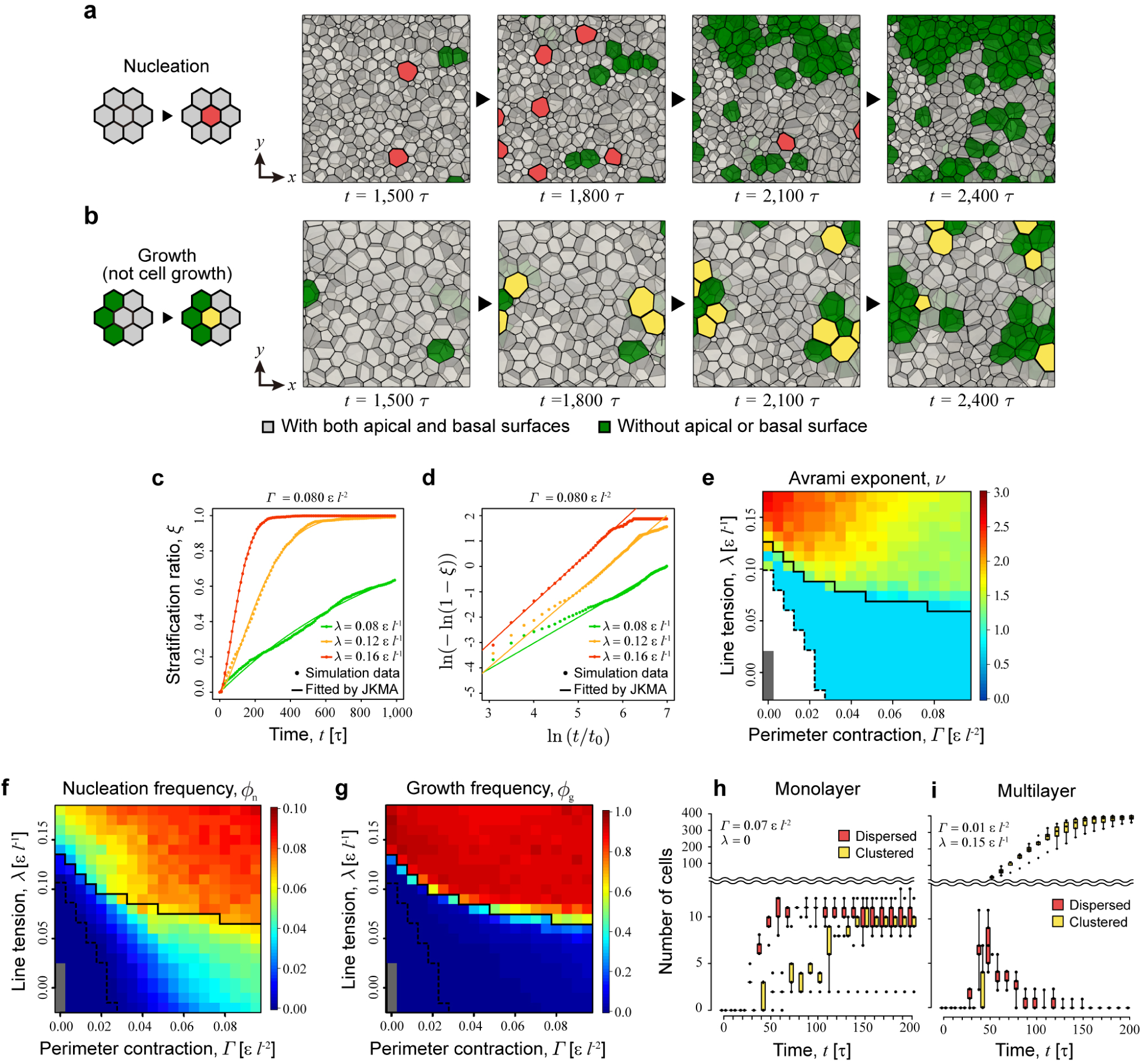
Three-dimensional vertex model reveals a nucleation-growth process driving epithelial stratification. **a**,**b** Schematic illustration (left) and corresponding simulation results (right) depicting nucleation and growth during stratification. Green cells represent those lacking apical, basal, or both surfaces. Red cells represent newly delaminated cells appearing in the monolayer, and yellow cells represent newly delaminated cells adjacent to existing ones. The parameters used were set to *Γ* = 0.085 *ε l*^-^*, λ* =0.07 *ε l*^-1^ (**a**) and *Γ* = 0.005 *ε l*^-2^*, λ* = 0.13 *ε l*^-1^ (**b**). **c**, **d** Linear and Avrami plots of the stratification ratio, *ξ*, as a function of time, *t*, respectively. Dotted lines indicate simulation data. Solid lines indicate lines fitted by the JMAK model. **e** Avrami exponent, *ν*, with respect to *Γ* and *λ*. The solid line indicates the boundary between the monolayer and multilayer structure. **f**, **g** Nucleation ratio, *ϕ*_n_, and growth ratio, *ϕ*_g_, with respect to apical perimeter contraction, *Γ*, and apical line tension, *λ*. **h**, **i** Temporal changes in the number of dispersed versus clustered delaminated cells under monolayer and multilayer conditions, respectively. In (**e**–**g**), solid and dotted lines indicate the boundaries among multilayer (*ξ >* 0.5), monolayer (0.5 ≥ *ξ* ≥ 0.1), and complete monolayer (0.1 > *ξ*) phases. In (**e**–**g**), the gray-colored regions indicate the collapsed state. In (**e**), the white-colored region denotes the monolayer state, for which the Avrami exponent was not calculated. The data in (**c**–**i**) indicate the average values calculated using five initial conditions. Box plots in (**h**, **i**): center: median; bounds, 25th and 75th percentiles; whiskers extend 1.5 times the interquartile range from the 25th and 75th percentiles; outliers are represented by dots.

Notably, the region with low *Γ* and high *λ* showed the closest Avrami exponent (*ν* ≈ 2.29) to that observed in the MDCK cell culture, corresponding to a region with a similar number of layers (Fig. 1d). These alignments between the simulations and experiments suggest that our model successfully recapitulates the kinetic process of epithelial stratification at single-cell resolution.

To investigate the nucleation-growth process, we calculated the ratios of the numbers of nucleation and growth, denoted by *ϕ*_n_ and *ϕ*_g_, respectively (detailed in Materials and Methods), and examined their dependencies on apical contractility: *Γ* and *λ*. Both *ϕ*_n_ and *ϕ*_g_ increased with *Γ* and *λ* (Fig. 3f, g). Importantly, *ϕ*_n_ alone did not distinguish between monolayer and multilayer structures: a monolayer formed at low *ϕ*_n_ (≲ 0.06) when *Γ* < 0.04, while a multilayer structure formed at high *ϕ*_n_ (≳ 0.02) when *Γ* > 0.04 (Fig. 3f). In contrast, *ϕ*_g_ differentiated these structures effectively: monolayer and multilayer domains corresponded to lower (≲ 0.1) and higher (≳ 0.8) *ϕ*_g_ values, respectively, across the entire range of *Γ* (Fig. 3g). Moreover, we examined stratification processes and found that, under monolayer conditions, nucleation initially increased, but growth did not occur (Fig. 3h). In contrast, under multilayer conditions, nucleation initially increased during the early stage of stratification, followed by an increase in growth (Fig. 3i), consistent with observations in MDCK cell culture (Fig. 1i). These findings indicate that the presence of domain growth after its nucleation determines the occurrence of the monolayer-to-multilayer transition.

These results demonstrate that, in both the MDCK cell culture and the simulations based on foam mechanics, a small portion of cells initially delaminates individually, and the stratified domains gradually expand through domain growth (Fig. 1g, h; Fig. 3a, b), rather than the majority of cells delaminating simultaneously across the entire epithelium. Consequently, the monolayer-to-multilayer transition is driven by a nucleation-growth process rather than spinodal decomposition.

### Foam-geometric instability induces the nucleation-growth process

To understand the mechanisms inducing nucleation and domain growth (i.e., single cell delamination), we developed a theoretical model based on our previous theory^30^. In this model, we focus on the deformations of a single cell and its adjacent cells within a planar monolayer and derive an energy function for a delaminating cell (see Materials and Methods). This model simplifies the situation by approximating Equation 3 using a mean-field approach, in which the heterogeneity of surrounding cell shapes is disregarded.

We examined the dependence of stable cell shapes on apical line tension (*λ*), planar cell density (*ρ*), and the number of adjacent cells (*n*) by analyzing energy landscapes. Here, to describe the shape of a delaminating cell, we introduced the cell taper, *d* (Fig. 4a), which satisfies −1 < *d* < 1 when the cell is embedded in a monolayer and becomes −1 or 1 when it delaminates in the basal or apical direction. Stable cell states were determined by identifying *d* which minimizes the energy function, denoted by *d*\*. We found that as *λ*, *ρ*, and *n* were varied, *d*\* shifted from −1 < *d*\* < 1 to *d*\* = −1 and/or *d*\* = 1, indicating the transition from an embedded to a delaminated state (Fig. 4b–d). Specifically, *d*\* bifurcated from −1 < *d*\* < 1 to *d*\* = −1 and 1 as a function of *ρ* and *n*, (Fig. 4c, d). These results indicate that mechanical instability is inherent in the foam-like epithelial geometry; either an increase in *λ*, an increase in *ρ*, or a decrease in *n* leads to cell delamination (Fig. 4e). To experimentally validate this, we calculated the probability distribution of *d* in MDCK cell culture (Fig. 4f). We found a trimodal probability distribution around *d* = −1, 0, and 1, with two distinct gaps between *d* = - 1 and 0 and between *d* = 0 and 1. This finding indicates that cells adopt either the embedded or delaminated state rather than staying in their intermediate state, implying that cells that lose the stability of the embedded state will immediately transition to the delaminated state. This observation supports the hypothesis that an inherent instability leads cells to either an embedded or detached state.

**Fig. 4:**
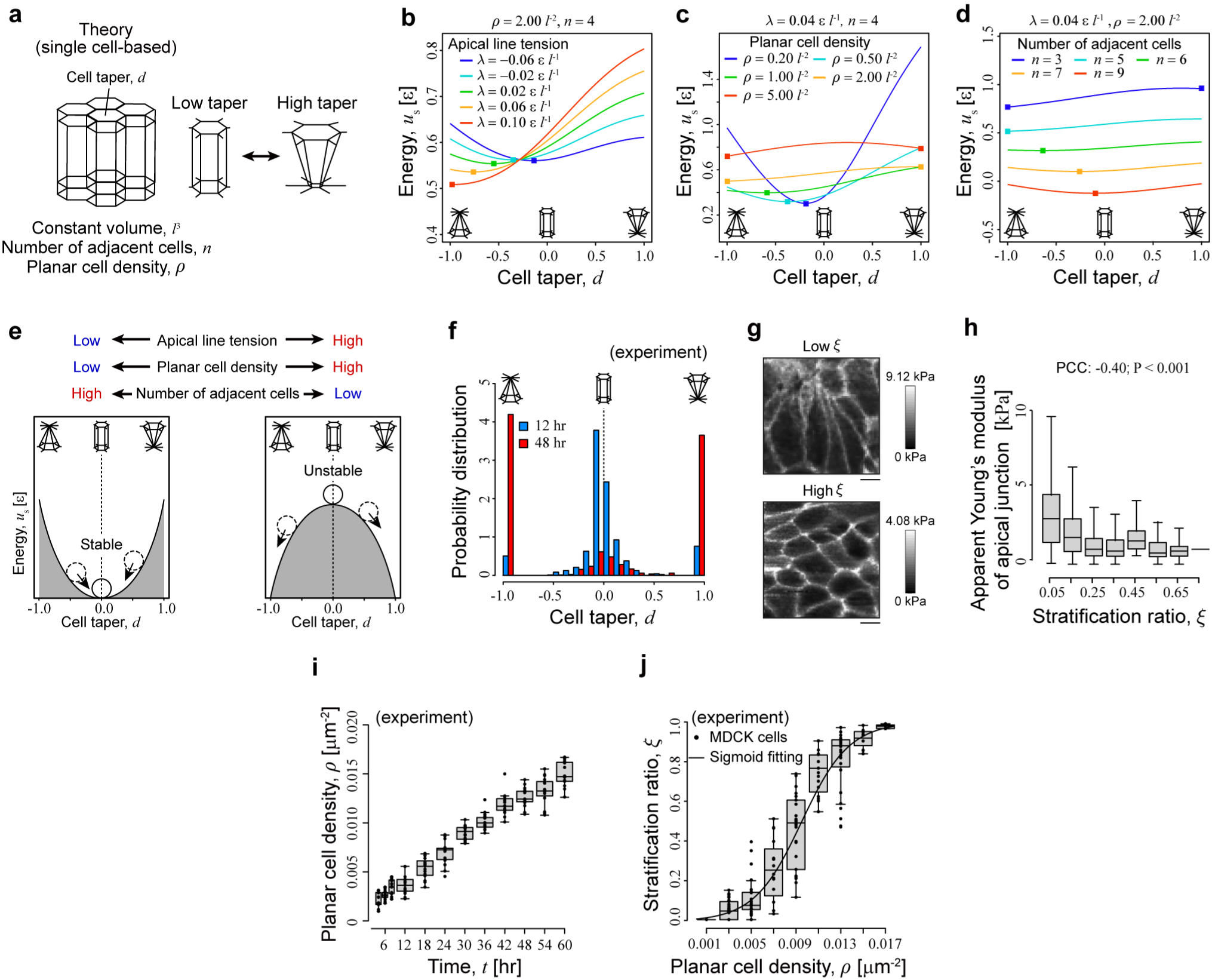
Foam-geometric instability induces epithelial stratification with scutoid formation. **a** Theoretical model describing the epithelial sheet 3D foam geometry. A single center cell is embedded within a sheet, modeled as a regular frustum, with its geometry uniquely determined by the number of adjacent cells, *n*, planar cell density, *ρ*, and cell taper, *d*. **b**–**d** Energy landscapes as a function of *d*, with respect to apical line tension *λ*, *ρ*, and *n*, respectively, derived from the theoretical model. Squares indicate energy minima. **e** Schematic illustration of mechanical instability driving the monolayer-to-multilayer transition. The state of the embedded cell switches between stable and unstable depending on *λ*, *ρ*, and *n*. **f** Probability distribution of cell taper calculated from experimental MDCK data at 12 (blue) and 48 (red) hours of culture (N ≥ 5). **g** Apparent Young’s modulus maps with high and low *ξ*. **h** Apparent Young’s modulus of apical junction as a function of *ξ*, obtained from AFM measurements (N = 19). Scale bar: 10 µm. **i**, **j** Stratification ratio, *ξ*, as a function of time, *t*, and *ρ*, respectively (N ≥ 15). The solid line in (**j**) represents a fitted sigmoid function. Plots in (**h**-**j**) include data from more than 15 images. Box plots in (**h**-**j**): center, median; bounds, 25th and 75th percentiles; whiskers extend 1.5 times the interquartile range from the 25th and 75th percentiles; outliers are represented by dots; ***P < 0.001 (two-sided Welch’s t-test). In (**h**), individual data points are not shown due to the presence of 10,000 points. In (**h**), the Pearson correlation coefficient (PCC) between the apparent Young’s modulus and *ξ* was calculated; P-values were obtained using Student’s t-test for non-correlation.

While the dependences on *λ*, *ρ*, and *n* corresponded to the previous reports showing the contributors to cell delamination, including apical junctional forces^19–24^, planar cell density^25–27^, and cell geometry^28–31^, the key driving force of epithelial stratification in MDCK cell culture (Fig. 1b) remained unclear. To clarify this, we examined each contributor. First, to determine whether the average value of the apical line tension *λ* drives epithelial stratification, we specifically measured the apparent Young’s moduli at apical junctions using AFM (Fig. 4g), which reflects the average value of junctional tensions^50^. We examined the correlations between the modulus and the stratification ratio, *ξ* and found that the apical junctional modulus decreased significantly with increasing *ξ* (Fig. 4h), which is in stark contrast to the theoretical prediction (Fig. 4b). These results suggest that apical junctional tension does not act as a key driving force behind stratification in MDCK cell culture.

Second, to determine whether *ρ* drives epithelial stratification, we calculated the planar cell density, *ρ*, in MDCK cell culture, defined as the inverse of the average apical and basal surface area per cell, normalized by cell volume (Fig. 4i; detailed in Materials and Methods). We observed a monotonic increase in *ρ* over time. Moreover, by correlating *ξ* with *ρ* over time, we assessed the dependence of *ξ* on *ρ* (Fig. 4j) and found that *ξ* increased with *ρ* in a switch-like manner, well-fitted by a sigmoidal curve. These results suggest that both nucleation and growth induced by the increase in cell density play critical roles in driving epithelial stratification. This finding aligns with previous studies highlighting the influence of cell density on individual cell delamination^25–27^ as well as our analytical calculations (Fig. 4c). Thus, planar cell density, rather than apical junctional tension, is the key driving force behind epithelial stratification in MDCK cell culture.

### Cell delamination proceeds via scutoid formation

To investigate the influence of *n* on stratification, we observed the changes in cell geometries during the delamination process in both simulations and experiments (Fig. 5a–i). During stratification, each cell gradually reduced the number of adjacent cells and lost either its apical or basal side to delaminate from the monolayer (Fig. 5a, c), which aligns with our theory (Fig. 4d). These processes can be represented by topological networks of cell-cell contacts (Fig. 5b, d), where cells are described as nodes and contacts as edges. Specifically, the contact edges were categorized into three types: (1) contacts through lateral boundary faces that connect to both the apical and basal surfaces (black lines); (2) contacts that connect only to the apical surface (red lines); and (3) contacts that connect only to the basal surface (blue lines). In both experimental and simulation results, although contact edges on the apical side were largely preserved, those on the basal side changed dynamically; contacts between delaminating cells and their neighboring cells were replaced by new contacts formed between neighboring cells on the basal side.

**Fig. 5:**
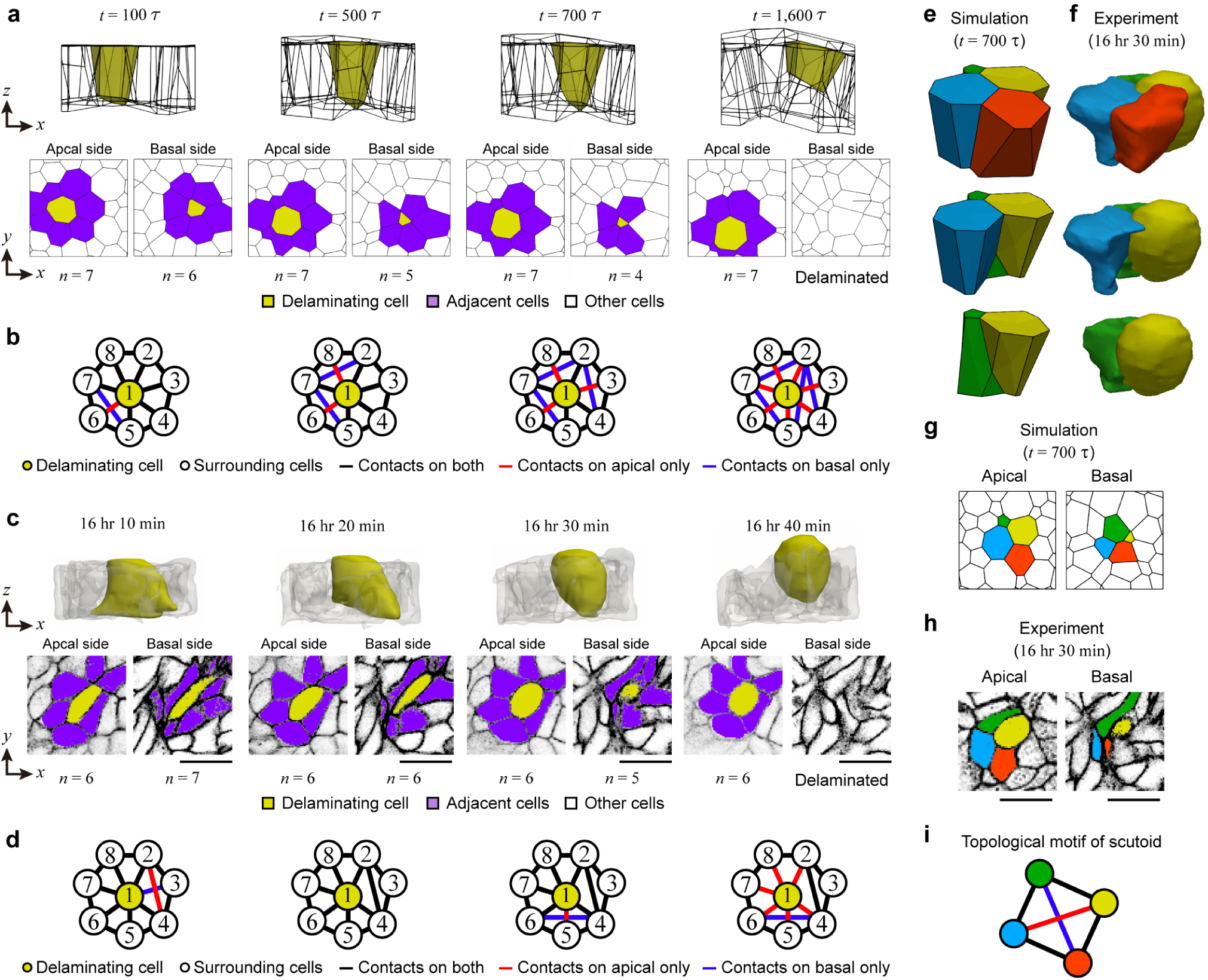
Cell delamination proceeds with formation of a scutoid. **a**, **c** Delamination processes in the 3D vertex model (**a**) and MDCK cell culture (**c**). The top images depict 3D views, and the bottom images show cross-sectional views of the apical and basal surfaces, with the number of cells adjacent to delaminating cells, *n*, indicated below. In the top and bottom panels, yellow highlights the cell delaminating toward the apical side as cells lose basal surfaces. In the bottom panels, purple marks cells adjacent to delaminating cells. Scale bar: 10 µm in (**c**). **b, d** Topological networks of cell-cell contacts on the apical and basal surfaces. Yellow and white nodes represent delaminating and neighboring cells, respectively. Black, red, and blue edges indicate cell-cell contacts present on both the apical and basal sides, the apical side only, and the basal side only. **e**–**h** Scutoid formation during delamination in the 3D vertex model (**e**, **g**) and MDCK cell culture (**f**, **h**). **e**, **f** 3D geometries of cells forming scutoids. The yellow cells correspond to those in (**a**, **b**) and (**c**, **d**), respectively. **g**, **h** Apical and basal shapes of the cells shown in (**e**) and (**f**), respectively. **i** Topological motif characteristic of scutoid geometry. This motif corresponds to the cell geometry shown in (**e**–**h**). Yellow, red, blue, and green nodes in (**i**) represent the respective cells in (**e**–**h**). Scale bar: 10 µm. In (**a**, **b**, **e**, and **g**), the parameters used were set to *Γ* = 0.085 *ε l*^-2^ and *λ* =0.07 *ε l*^-1^.

Interestingly, during the delamination process, we found that cells formed a scutoid geometry ^51–53^, which emerged when cell-cell contacts differed between the apical and basal sides. In such cases, cell-cell contacts switched along the lateral surfaces between neighboring cells (Fig. 5e, f). Specifically, the blue and yellow cells made contact on the apical side, whereas the red and green cells made contact on the basal side (Fig. 5g, h). The scutoid can be characterized by a specific topological motif within a local network of cell-cell contacts (Fig. 5i). In this motif, four cells, represented as nodes, form a quadrilateral contact topology, within which each cell exhibits different edge connections on the apical and basal sides, thereby adopting a scutoid geometry. Such motifs can be identified in the topological contact networks observed during stratification (Fig. 5b, d). For example, in the numerical simulation, cells 1 and 3 formed a contact, while cells 2 and 4 formed contacts at *t* = 500 *τ*, indicating that cells 1–4 collectively form scutoids (Fig. 5b).

In epithelia of a certain thickness, different cell-cell contacts often arise between the apical and basal surfaces. When cell rearrangements occur within such epithelial planes, cells transiently adopt scutoid geometries, which enable topological transitions through the exchange of cell-cell contacts within the plane. Consistent with this phenomenon, most scutoids were observed to appear transiently during cell delamination (Fig. 5b, d). Scutoid formation is associated with an increase in the total number of cell-cell contacts across the tissue, facilitating the rearrangement of neighboring contacts required for delamination. Ultimately, delaminating cells reduce the number of their cell-cell contacts on either the apical or basal side, allowing them to exit the epithelial layer.

These results demonstrate that mechanical instability inherent in the foam-like epithelial geometry induces nucleation and domain growth. This instability can be triggered by an increase in apical junctional tension or planar cell density. During cell delamination, cells gradually lose either their apical or basal side while forming a scutoid geometry to reduce the number of adjacent cells, which further contributes to the onset of instability. The accumulation of cell delaminations leads to the phase transition of the epithelial structure from a monolayer to a multilayer.

### Nucleation-growth concept applies to embryogenesis and carcinogenesis

To determine whether the nucleation-growth concept applies to embryogenesis and carcinogenesis, we examined the epithelial stratification processes during mouse skin development and malignant transformation of mouse intestinal cancer organoids (Fig. 6). Skin development involves epithelial stratification, during which the epidermal monolayer transitions into a multilayer structure. Initially, basal cells delaminate and give rise to an intermediate suprabasal layer composed of poorly differentiated cells. This layer subsequently differentiates into committed periderm cells that cover the epidermis^12,54,55^ (Fig. 6a). This stratification initiates at the tips of the limbs and tail and gradually extends across the body^54^. However, the mechanism underlying the spread of the stratification domain remains poorly understood. To address this issue, we analyzed the dorsal skin of mouse embryos (Fig. 6a; see Materials and Methods), identifying three distinct structural states from embryonic day E9.5 to E13.5: near monolayer, mixed layer, and multilayer (Fig. 6b). Spatial analysis of delaminated cells in the epidermis revealed two patterns: dispersed and clustered arrangements (Fig. 6c). These observations indicate that epidermal stratification proceeds through both nucleation and growth, rather than through continuous growth of multilayered domains.

**Fig. 6:**
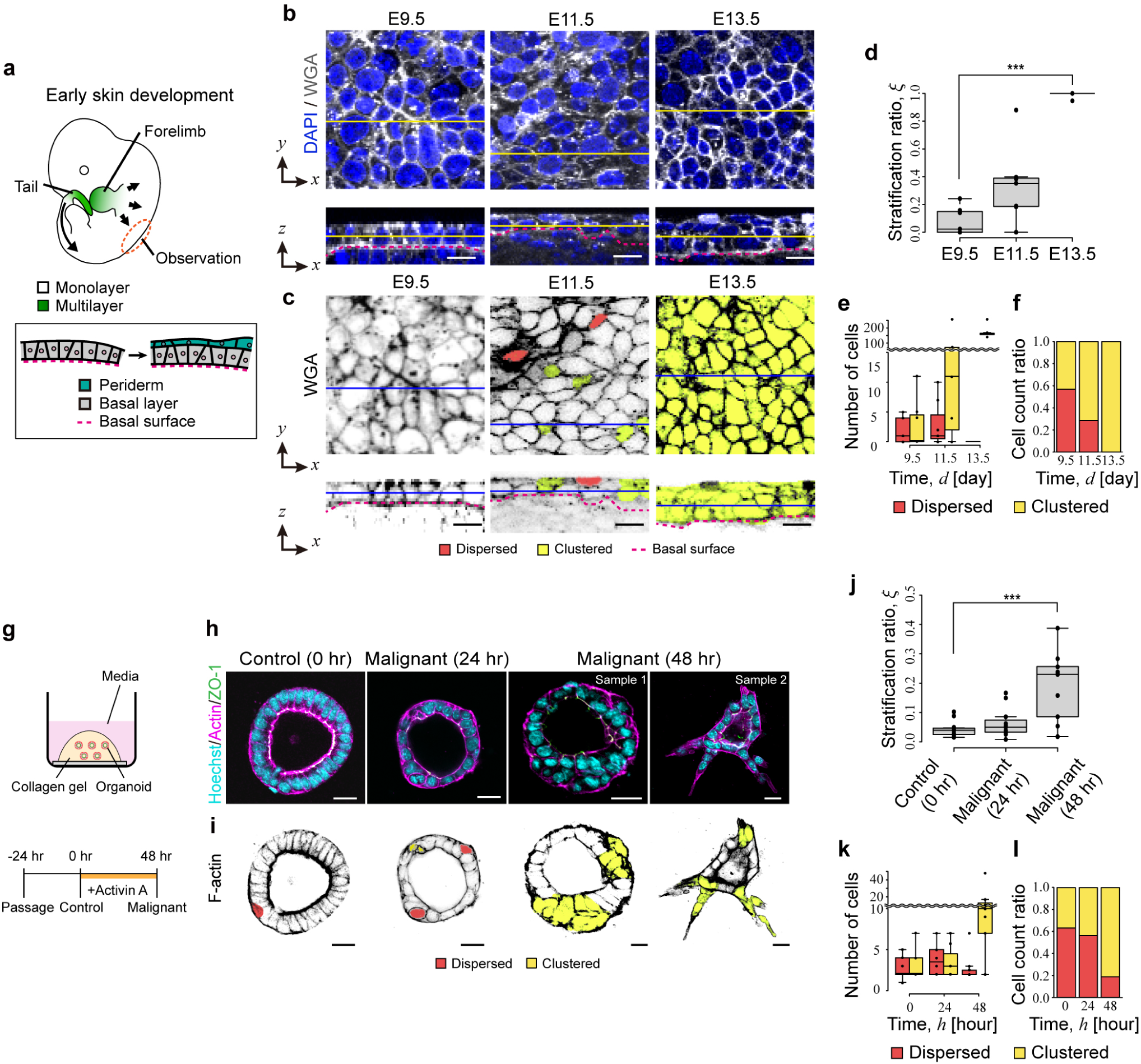
Nucleation-growth concept applies to embryogenesis and carcinogenesis. **a**-**f** Data for mouse skin development. **a** Schematic illustration of epidermal stratification progression in the entire mouse embryo. **b**, **c** Fluorescent section images of mouse skin and modified images highlighting delaminated cells. The skin is a monolayer in E9.5, mixed layer in E11.5, and multilayer structure in E13.5. Red and yellow cells represent dispersed and clustered cells. The top panel shows the xy-view, and the bottom panel shows the *xz*-view. Yellow and blue lines indicate cross sections. The dotted lines mark the basal surfaces. Scale bar: 10 µm. In (**b**), epidermises were stained with wheat germ agglutinin (WGA) antibody and DAPI. **d** Stratification ratio, *ξ*, in each stage. **e** Temporal changes in the number of dispersed versus clustered delaminated cells. **f** Temporal changes in the ratio of dispersed and clustered delaminated cells. **g**-**l** Data for malignant transformation of intestinal cancer organoid. **g** Schematic illustration of cancer organoid culture and malignant phenotype induction. **h**, **i** Fluorescent section images and modified images highlighting delaminated cells, respectively. In (**h**), organoids were stained with a ZO-1 antibody, phalloidin, and Hoechst. Scale bar indicates 20 µm. In (**i**), red and yellow cells represent dispersed and clustered cells. **j** Stratification ratio, *ξ*, under control and malignant induction. **k** Temporal changes in the number of dispersed versus clustered delaminated cells. **l** Temporal changes in the ratio of dispersed and clustered delaminated cells. In (**c**, **i**), dispersed and clustered delaminated cells are colored red and yellow, respectively. Plots in (**d**–**f**, **j**–**l**) include data from more than five images. Box plots in (**d**–**f**, **j**–**l**): center: median; bounds, 25th and 75th percentiles; whiskers, min and max; ***P < 0.001 (two-sided Welch’s t-test).

To clarify whether epidermal stratification proceeds via a nucleation and growth process, we quantified the layer structures of the epidermis. The stratification ratio increased from approximately 0.08 to 1.0 during the monolayer-to-multilayer transition (Fig. 6d). Moreover, to understand the time evolution of the nucleation and growth process during skin development, we determined whether the cells were dispersed or clustered. Dispersed cells increased during the early stages of stratification, followed by an increase in clustered cells, similar to the behavior observed in MDCK cells (Fig. 6e, f). These findings suggest that the concept of a phase transition, involving nucleation and growth, is applicable to skin development.

To investigate whether this concept also applies to cancer progression, we studied early-stage cancer cell invasion, where precancerous epithelial cells undergo abnormal stratification leading to invasion. For this purpose, we used key cancer-driving mutations in mouse intestinal cancer organoids: Apc (A), Kras (K), Tgfrbr2 (T), Trp53 (P), and Fbxw7 (F) (see Materials and Methods) ^56^. A previous study demonstrated that exposure to activin A in culture (Fig. 6g) induces malignant transformation in these organoids, mimicking *in vivo* cancerous behaviors^57^. Specifically, organoids cultured without activin A formed spherical cysts, whereas those exposed to activin A displayed malignant morphologies, such as heterogenous spherical or protruded, star-like structures (Fig. 6h). At 0 hours, organoids exhibited a monolayer structure with a few dispersed delaminated cells. By 48 hours post-transformation, these organoids transitioned to a multilayered structure with clustered delaminated cells (Fig. 6i).

Consistent with the observations, stratification ratios increased from approximately 0.02 to 0.2 during this transition (Fig. 6j). Moreover, to understand the time evolution of the nucleation and growth process during malignant transformation, we determined whether cells were dispersed or clustered (Fig. 6k, l). Dispersed cells increased during the early stages of stratification, followed by an increase in clustered cells, similar to the behavior observed in MDCK cells. These results suggest the analogy between the phase transition involving nucleation and growth and malignant transformation and demonstrate that the nucleation-growth concept applies beyond only MDCK cell culture to embryogenesis and carcinogenesis.

## Discussion

In this study, we elucidated the physical mechanism underlying epithelial stratification, focusing on the transition from a monolayer to a multilayer structure (Fig. 7). We discovered that this is a first-order phase transition, involving nucleation and growth. The growth rate of delaminated domains plays a more decisive role in the monolayer-to-multilayer transition than nucleation. The elementary process of both nucleation and growth, namely single-cell delamination, is induced by the mechanical instability inherent in foam-like epithelial geometry. This instability can be evoked by an increase in either apical junctional tension or planar cell density, regulated by various cell behaviors such as actomyosin contractility^19–24^, cadherin adhesion^58–60^, and cell proliferation^25–27^. The instability can also be triggered by a decrease in the number of cells adjacent to a delaminating cell, mediated by the formation of cell scutoid geometry. Overall, these findings demonstrate that epithelial stratification can be understood as a phase transition governed by foam mechanics. This analogy could be applied to general epithelial stratification, including those occurring during embryogenesis and carcinogenesis.

**Fig. 7:**
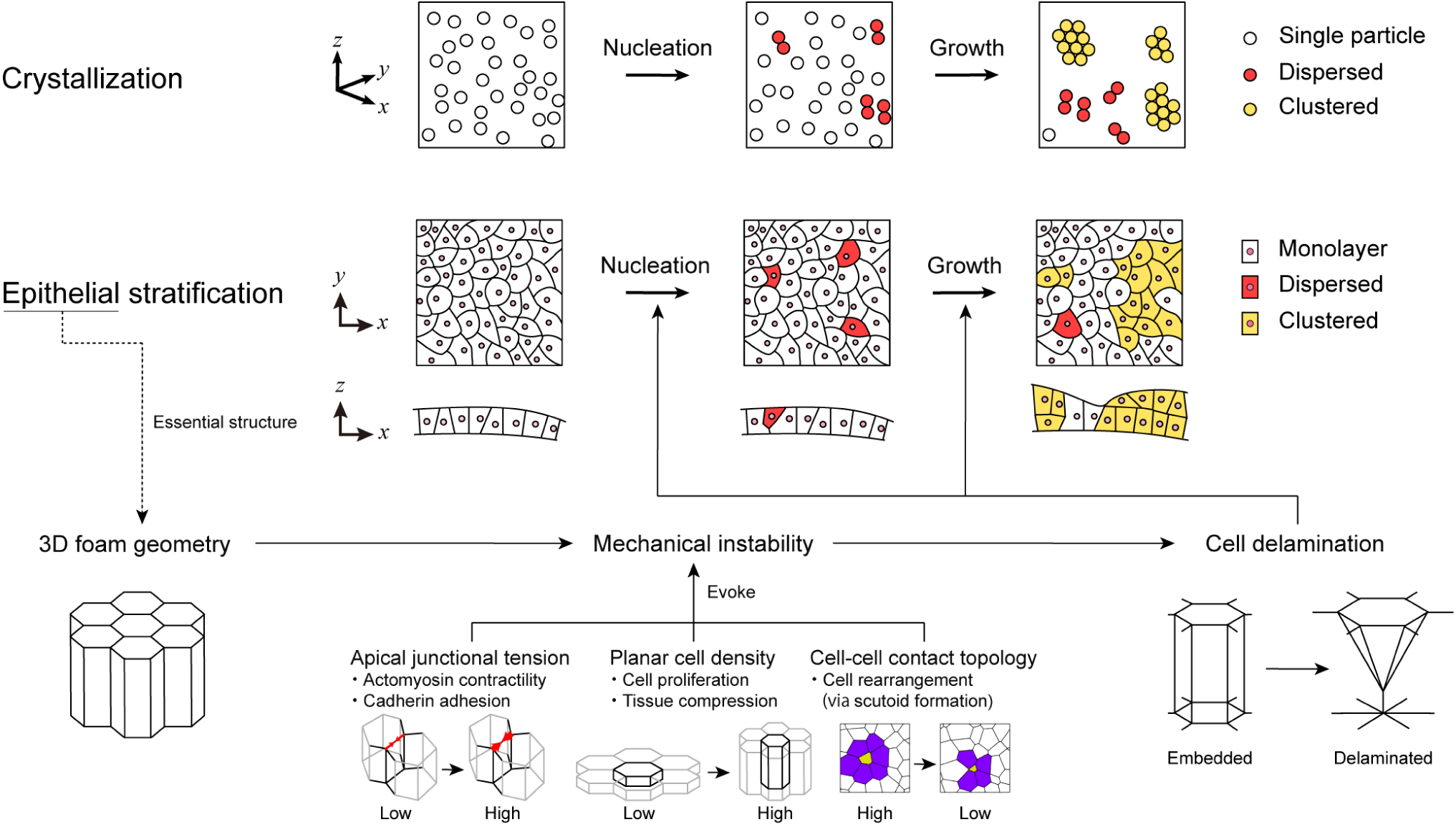
Epithelium stratifies via nucleation and growth induced by foam-geometric instability. Epithelial stratification can be understood as a phase transition, such as crystallization, that involves a nucleation-growth process. In this framework, stratification represents a structural transition from a monolayer to a multilayer configuration. The fundamental process driving this transition is cell delamination, which arises from mechanical instability inherent in the 3D geometry of the epithelial sheet. This instability can be evoked by various factors, including increased apical junctional tension due to actomyosin contractility and cadherin-mediated adhesion, increased planar cell density resulting from cell proliferation and tissue compression, and changes in cell-cell contact topology caused by rearrangements associated with scutoid formation.

The process of nucleation and growth is a well-established concept in solid matter physics. A pioneering study demonstrated that, in epithelial cell clustering, the transition from dispersion to aggregated states proceeds via a phase transition driven by nucleation and growth processes^61^. Nevertheless, to the best of our knowledge, this is the first report of its application in the context of epithelial stratification. Traditionally, crystallization of materials, such as water and metals, is driven by nucleation and growth, typically induced by an increase in material density^62^. Our research demonstrated that epithelial stratification shares these characteristics (Fig. 1–3), highlighting their similarities with nucleation-and-growth processes in a wide range of biological phenomena, including embryogenesis and carcinogenesis (Fig. 6). Based on this analogy, we can apply methods used in the physical understanding of nucleation and growth to investigate epithelial stratification. For instance, using the JMAK model^42^, we evaluated whether nucleation or growth drives each stratification process. Specifically, frequent growth was observed in MDCK cell culture (Fig. 1d, e). This analysis will be helpful in exploring cell behaviors in the developmental mechanisms of various epithelia^1–9^, teratogenic diseases caused by stratification failure, such as lissencephaly and neuronal migration disorders^63^, and malignant transformation initiated by stratification^13^. Therefore, there is potential applicability of existing methods for nucleation and growth to deepen the understanding of a wide range of phenomena involving epithelial stratification, including embryogenesis and carcinogenesis.

Other physical behaviors may also contribute to the transition from a monolayer to a multilayer structure in epithelia. One possibility is active cell migration^64^, which has been observed in wound healing^65^ and morphogenetic processes^66^. Cells actively move and rearrange in response to internal or external cues, potentially leading to local overcrowding and cell delamination. Another candidate is cell sorting^67^, driven by differential adhesion or cortical tension, which can cause spatial segregation of cell populations and potentially result in multilayered structures, particularly when combined with variations in cell type or mechanical properties. Moreover, several biological behaviors may also contribute to this transition, such as the orientation of cell division directions^14–16^, differences in proliferation rates^15^, and signaling patterns^68,69^. These alternative behaviors highlight the complexity of epithelial morphogenesis and suggest that multiple, possibly concurrent, mechanisms may underlie stratification depending on the biological context.

While our model revealed that an increase in planar cell density facilitates epithelial stratification, high density alone does not always lead to stratification. Our model also predicted that whether stratification occurs depends not only on cell density but also on other parameters, such as apical junctional tension. Consistently, the *Drosophila* imaginal disc epithelium, a pseudostratified monolayer, maintains its structure despite increased proliferation and crowding^70^. Similarly, larval salivary glands remain as a monolayer even though their cells are large and polyploid^71^. These observations suggest that additional mechanical factors, such as apical junctional tension, cell–ECM interactions, and cell-nuclear interactions, play critical roles in determining epithelial layer architecture. The deformability of individual cells, as observed in pseudostratified tissues, may also be a key determinant. Models that incorporate more flexible representations of cell shape^72,73^ may help address these complexities and further refine our understanding of stratification mechanisms.

Recent studies have revealed that scutoids emerge frequently in epithelia with high cell density or high curvature or after proliferation events^51–53^. Our model and experiments revealed that scutoids also appear during epithelial stratification. However, scutoid formation does not always induce stratification. Consistently, tissues such as the salivary gland^51^ and sea star blastulae^53^ exhibit scutoid-rich architectures while maintaining a monolayer. These findings suggest that scutoids represent an intermediate state of cell rearrangement, enabling topological transitions through the exchange of cell-cell contacts within the epithelial plane.

While this study provides a conceptual understanding of epithelial stratification, it is important to acknowledge its limitations. Through our experiments, the nucleation-growth process was observed in MDCK cell cultures, skin development, and cancer cell invasion, suggesting the applicability of this concept to a wide range of epithelial stratification. On the other hand, the contribution of mechanical instability to cell stratification in biological systems remains only partially understood. The proposed theoretical model, exhibiting mechanical instability, showed good agreement with perturbative experiments in MDCK cell cultures as well as previous studies, thereby supporting its applicability. Nevertheless, clarifying its role in embryogenesis and carcinogenesis will require further investigation. Additionally, our model simplifies the problem of energy minimization for a fixed number of cell populations with uniform properties. The model also adopts stress-free boundary conditions; however, *in vivo* epithelial tissues are subject to various external forces, including confinement by surrounding tissues and interactions with the substrate and ECM. These factors may significantly influence cell delamination and stratification^74,75^. This simplification allows clarification of the fundamental understanding, whereas the roles of more complex cell behaviors remain unclear, such as cell proliferation, cell motility, and the diversity in cell properties caused by differentiation, all of which could play significant roles in regulating stratification in broader biological scenarios^14–16,69,71,76–78^. As such, the relationship between our experiments and simulations remains primarily qualitative. This study marks the beginning of a multiscale physical exploration aimed at understanding the entire stratification process. Incorporation of these additional cellular behaviors in future research could lead to a more quantitative understanding of critical biological processes such as embryonic development and cancer progression.

## Methods

### MDCK cell line

Madin-Darby canine kidney (MDCK, type I) cells stably expressing mCherry-CAAX were provided by Dr. Kajiwara (Osaka University, Osaka). The cells were cultured in Dulbecco’s Modified Eagle’s medium (DMEM, Nacalai Tesque) supplemented with 10% fetal bovine serum (FBS, Gibco) and 1% penicillin-streptomycin (Wako). Cultures were maintained in a 5% CO_2_ incubator at 37°C.

### MDCK cell culture to observe epithelial stratification

A 35-mm glass-bottom imaging dish (Matsunami) was coated with 30 µL of Cellmatrix Type I-A (Nitta Gelatin). A cell suspension containing 1.8 × 10^5^ cells in 250 µL of DMEM (Nacalai Tesque) supplemented with 10% FBS (Gibco) was added to the dish and incubated. After 4 hours, 2 mL of the medium was added. The dish was incubated until 60 hours after addition of the cell suspension.

### Immunostaining of MDCK epithelium

Cells were fixed in 4% paraformaldehyde (PFA) (Fujifilm Wako) in phosphate-buffered saline (PBS) for 30 minutes at room temperature (RT), followed by two washes with PBS. The cells were permeabilized with 0.5% Triton-X in PBS for 2 minutes. After three washes with PBS, the samples were blocked with 4% bovine serum albumin (BSA)/PBS for 10 minutes and incubated for 1.5 hours with anti-ZO-1 (rabbit/1:100 dilution, 61-7300, Invitrogen). After washing with PBS, the samples were incubated for 1.5 hours with Goat anti-Rabbit IgG 488 H&L antibody (1:100 dilution, ab150077, Abcam), Phalloidin-iFluor 647 Conjugate (1:1000 dilution, 20555, Cayman), and Hoechst (1:1000 dilution, 343-07961, Fujifilm Wako). Following three washes with PBS, the samples were mounted in 50% glycerol in PBS. Fluorescent images were acquired using a BZ-X810 microscope (Keyence).

### Observation and quantification of MDCK epithelium

Z-stack imaging was performed at each time point using a BZ-X810 microscope (Keyence) equipped with a 60× (NA 0.45) objective and a LMS800 microscope (Zeiss) equipped with a 25× (NA 0.8) objective. Images were captured using the 2× zoom function with a step size of 0.4–2 µm. Five images were obtained from the center to the periphery of the dish at each time point.

The stratification ratio, *ξ*, was calculated using Equation 1. For this calculation, cells were categorized into four types: cells with both apical and basal surfaces, cells with either apical or basal surfaces, and cells with neither apical nor basal surfaces (Fig. 1c). The presence of apical and basal surfaces was manually identified using an Advanced Observation Module (BZH4XD, Keyence), based on the mCherry-CAAX intensity. *ξ* was calculated at every time point and fitted with the JMAK model in Equation 2 using a nonlinear least squares method to obtain the parameters *K*, *n*, and *t*_0_. Moreover, delaminated cells were classified as dispersed and clustered based on the presence of adjacent delaminated cells using Fiji software.

Planar cell density, *ρ*, was determined by counting cells based on the mCherry-CAAX intensity. The number of cells was calculated as *N*_c_ + *N*_e_⁄2, where *N*_c_ is the number of single cells fully located within an image and *N*_e_ is the number of cells partially crossing the image edge. *ρ* was calculated by dividing the number of cells by the total area of the image. Moreover, the correlation between *ξ* and *ρ* was determined by calculating *ξ* and *ρ* for individual images.

### Observation and visualization of single MDCK cell shapes

Time-lapse imaging was performed using an FV 3000 confocal microscope (Olympus) equipped with a GaAsP detector (FV31-HSD, Evident) and a dichroic mirror (U-FBNA, Olympus) in a humidified 5% CO_2_ atmosphere at 37°C. Images were captured with a step size of 0.43 for 6 hours, at 10-minute intervals. Using the mCherry-CAAX intensity, individual cells were manually masked using Fiji and Drishti v. 3.2. The masked images were visualized using ParaView (Kitware Inc.) and Imaris viewer (Bitplane).

### Immunostaining and observation of mouse embryonic skin

To prepare mouse embryonic skin tissue samples, pregnant Jcl:ICR mice (CLEA Japan, Inc.) were euthanized by cervical dislocation, and embryos at E9.5, 11.5, and 13.5 were collected. The head, tail, limbs, and internal organs were removed, leaving the skin tissue, which was fixed with 4% PFA in PBS for 1 hour at 4°C. Following fixation, the tissues were washed with PBS and blocked with 1% BSA/PBS for 1 hour at 4°C. Nuclei were stained with DAPI (1:1000; Dojindo; D523), and plasma membranes were labeled with wheat germ agglutinin (WGA, 1:200; Invitrogen; W32466) for 2 hours at RT. The tissues were then washed with PBS and immersed in 60% 2,2’-thiodiethanol (FUJIFILM; 201-04193) in PBS for 1 hour at RT to enhance optical clarity. Fluorescence signals were captured with a TCS SP8X confocal microscope (Leica) with a 40× oil immersion objective lens (Leica, HC PL APO 40×/1.30 Oil CS2) and LAS X software (version [BETA] 3.5.7.23723).

### Quantification of mouse embryonic skin

The stratification ratio, *ξ*, was calculated using Equation 1. For this calculation, we counted the total number of cells constituting epidermis in 3D stacked images based on both DAPI and WGA signals using LAS X. Delaminated cells in skin tissue were defined as cells without apical or basal surfaces, and the number of cells was manually counted. Basal surfaces were defined as having high area fluorescence intensity of the WGA signal. Seven samples were randomly selected for each stage.

### Intestinal cancer cell line

Intestinal cancer cells, provided by Dr. Oshima (Kanazawa University, Kanazawa), were derived from mouse somatic cells that had been transduced with cancer mutations, as previously described^56^. These cells were originally harvested from small intestinal tumors in mouse models that carried a combination of key colorectal cancer driver mutations — Apc (A), Kras (K), Tgfrbr2 (T), Trp53 (P), and Fbxw7 (F) — in their intestinal epithelial cells. These cells reflect *in vivo* malignant behaviors, serving as a realistic model to study cancer progression and transformation.

### Intestinal cancer organoid culture

Intestinal cancer organoids were prepared using a previously described method^79^. Intestinal cancer cells were enzymatically or mechanically dissociated into either single cells or small aggregates and then cultured at 37°C in a 48-well dish (Corning) embedded in 30 µL of Collagen gel (Corning) surrounded by 300 µL of a media cocktail for organoid culture. The media contained Advanced DMEM/F-12 medium (Gibco), supplemented with 1% 100X GlutaMAX^TM^ (Gibco), 1% 100X N-2 (Gibco), 1% 1M HEPES buffer solution (Gibco), 1% penicillin/streptomycin solution (Wako, 100X), 2% 50× B-27 supplement (Gibco), and 0.1% 1 mM N-acetylcysteine (Sigma). Organoids were passaged every 2–3 days for maintenance by resuspending them while embedded in fresh Collagen gel with media in a new well of a culture dish.

### Malignant phenotype induction of intestinal cancer organoids

Malignant phenotypes were induced using a previously described method^57^. Intestinal organoids were passaged into a Collagen I matrix (Wako) using the media cocktail for organoid culture. After 24 hours of incubation at 37°C, the medium was replaced with fresh medium with an additional 20 ng mL^-1^ of the induction factor activin A (Cosmobio).

### Intestinal cancer organoid immunostaining and observation

Organoids were fixed by emersion in 4% PFA (Wako) at RT for 30 minutes. Samples were then washed three times in PBS before permeabilization and blocking. Permeabilization was a two-step process starting with emersion in 0.5% TritonX-100 (Fisher)/PBS for 30 minutes at 4°C. Next, samples were permeabilized in PBS containing 0.1% BSA (Wako), 0.2% TritonX-100, and 0.05% Tween20 (Wako) at RT for a minimum of 2 hours. The samples were blocked by incubation in 1% BSA solution for 1 hour at RT. They were then immunostained overnight at 4°C with nucleus (Cellstain Hoechst 33258, Wako, 1:300 dilution), F-actin (Phalloidin iFluor 647 Conjugate, Cayman, 1:500 dilution), and ZO-1 (ZO-1 Polyclonal Antibody, Invitrogen, 1:100) antibodies diluted in 1% BSA. Secondary antibody (Goat anti-Rabbit IgG H&L Alexa Fluor®488, Abcam, 1:200 dilution) incubation was performed under the same conditions as for primary antibodies after permeabilization in 0.1% BSA (Wako), 0.2% TritonX-100, and 0.05% Tween20 (Wako) buffer at RT for a minimum of 2 hours. Following immunofluorescence assays, fluorescent imaging of samples was performed using a confocal microscope (FV3000, Olympus) equipped with a 20× objective lens (20x-UPlanXApo, Olympus). Three-dimensional projections of images were reconstructed using ImageJ for analysis.

### Intestinal cancer organoid stratification ratio calculation

The stratification ratio, *ξ*, was calculated using Equation 1. The total number of cells constituting each organoid in 3D stacked images was counted based on both F-actin and Hoechst signals using Fiji software. Delaminated cells in the organoids were defined as cells lacking either apical or basal surface, similar to the definition used for MDCK cells. These cells were manually counted based on ZO-1 and phalloidin immunostaining. For the control group, five organoids were randomly selected from three separate culture dishes. For the activin-treated group, nine organoids were randomly selected from five dishes.

### 3D modeling of epithelial stratification

We employed a 3D vertex model to calculate the dynamics of multiple cells comprising an epithelial sheet in 3D space. The dimensions of the system box were defined as 0 ≤ *x* < *L*_*x*_, 0 ≤ *y* < *L*_*y*_, with z extending infinitely in both directions (−∞ < *z* < ∞) in the *xyz* coordinates. We applied periodic boundary conditions at *x* = 0, *L*_*x*_, *y* = 0, and *L*_*y*_. Within this box, an epithelial sheet, consisting of *N*_t_ cells, was arranged in the *xy* plane.

Each cell within the epithelium was modeled as a polyhedron, with shared vertices between adjacent cells. Therefore, the shape and arrangement of cells in the epithelium were described by the positions of the vertices of the polyhedrons and the topological connections among these vertices.

The epithelial polarity of each cell surface was defined based on this topology. The top and bottom surfaces of the sheet along the *z* axis were designated as the apical and basal surfaces, respectively, for each cell. Boundaries between adjacent cells were defined as lateral surfaces for each cell.

Cell movements are described by changes in the positions of the vertices. The time evolution of the position of the *i*th vertex, denoted by ***r***_*i*_, is described by

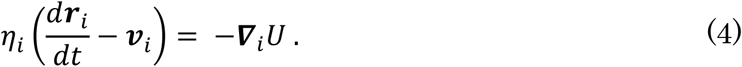

The left-hand side of Equation 5 represents the viscous force acting on the *i*th vertex. Here, *η*_*i*_ denotes the friction coefficient for this vertex, which is calculated as the sum of the friction coefficients from the adjacent cells, expressed as 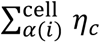, where *η*_*c*_ represents the friction coefficient between cells. The local velocity vector at the *i*th vertex, denoted by ***v***_*i*_, in Equation 5 is defined as the average velocity of the surrounding cells: 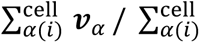, where ***v***_*α*_ is the velocity of the *α*th cell. ***v***_*α*_ is calculated as the average velocity of its constituent vertices. This formulation ensures the balance of the total viscous force within each cell. The right-hand side of Equation 5 represents the mechanical force acting on the *i*th vertex, which is derived from the effective energy of the system as described in Equation 3.

During epithelial stratification, cell configurations were rearranged in 3D space. In our model, this cell rearrangement is described by introducing [H]-[I] operations^34^. When individual edges in the network occasionally shrink to meet or retract from neighboring cells, the topological network is reconnected. These [H]-[I] operations differ from the well-known T1 transformation commonly used in 2D vertex models but express several changes in 3D cell configurations, including the rearrangement of cells within the plane of the epithelium, stacking of cells vertically on the plane, and hole formation within the epithelial sheet.

### Stress-free periodic boundary conditions

During the simulation, we ensured that the axial stress in the *xy* plane was approximately free by affinely deforming the system box. This deformation is described by

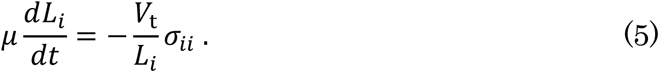

In this equation, the index *i* takes *x* and *y*, representing the respective axes. Here, *V*_t_ denotes the total volume of the cells, calculated as 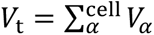. The constant *μ* is defined as the friction coefficient of the epithelial sheet to its surroundings. *σ*_*ii*_ represents the normal component of the stress tensor in the *i*th axis.

The stress tensor of the system, denoted by *σ*_*i*_*_j_*, is calculated by

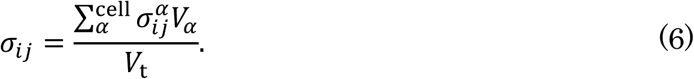

In this equation, 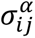 represents the stress tensor of the *α*th cell. We define 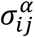 by

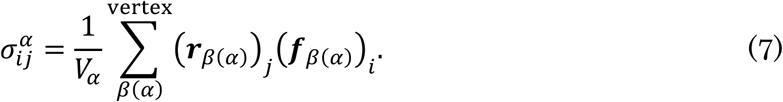

Here, ( )_*i*_ indicates the *i*th component of a vector. The vector ***r****_β_*_(*α*)_ is the distance vector from the center of the *α*th cell to the position of the *β*th vertex. ***f****_β_*_(*α*)_ represents the force acting on the *β*th vertex within the *α*th cell, given as ***f****_β_*_(*α*)_ = −*du*_*α*_/*d****r****_β_*, where *u*_*α*_ denotes the effective energy of the *α*th cell. The total effective energy *U,* described in Equation 3, is the sum of the effective energies of individual cells: 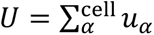, where the effective energy of each cell is given by

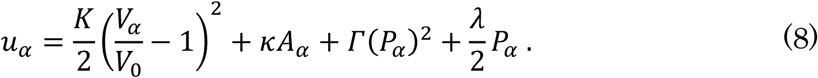

### Nondimensionalization and numerical simulations

Physical parameters were nondimensionalized by units: length, *l*, was scaled by 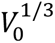, time, *τ*, by 12.5*η*_c_ *κ*^−1^, and energy, *ε*, by 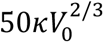. For the numerical integration of Equation 5, we employed the first order Euler’s method with a time step of Δ*t*, set to 0.005. Vertex velocities in Equation 5 were iteratively solved by convergent calculations to ensure that the mean residual error was below the threshold, *RE*_th_, set to 0.00001. Topological operations were applied to each edge and trigonal face when related edge lengths were under the threshold, Δ*l*_th_, set to 0.05, at each time interval of Δ*t*_r_, set to 1.0. Details of the numerical implementations are similar to those of our previous studies^34,35,47^.

To determine the local energy minimum for each parameter set, we calculated the relaxation process of vertex positions using Equation 5. The initial condition for these simulations was a disordered cell configuration within the monolayer. This condition was obtained by simulating tissue growth. This initial condition was created by proliferation from 100 cells to 401 cells under conditions that maintained the monolayer. In the numerical simulations, five disordered monolayer sheets created with different seed values were used as initial conditions. Numerical simulation data were averaged using these five data sets.

### Assessment of epithelial structures

The number of layers in the epithelium, *N*_l_, is defined as

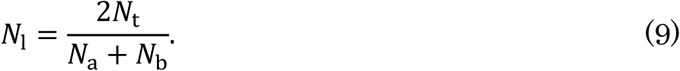

In this equation, *N*_a_ is the number of cells with an apical surface, and *N*_b_ is the number of cells with a basal surface. *N*_t_ is the number of total cells. This formula applies the trapezoidal formula and measures the average tissue height. This value represents the average number of layers in the tissue.

The nucleation ratio, *ϕ*_n_, and the growth ratio, *ϕ*_g_, are defined as

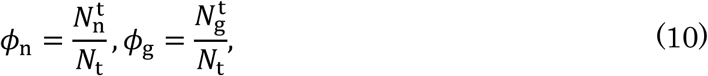

respectively. In this equation, 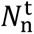 represents the total number of cells delaminated through the nucleation process, 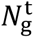 represents the total number of cells delaminated through the growth process, and *N*_t_ represents the total number of cells, including both delaminated and non-delaminated cells.

### Theoretical model of 3D foam geometry and mechanics

A theoretical model was developed to analyze single-cell delamination, as previously described^30^ For simplification, we assumed that the apical and basal surfaces were constrained in the plane and considered only the movements of a single center cell and its first neighbors within the sheet while keeping the other neighboring cells fixed in position. Under this approximation, the in-plane cell density *ρ* implicitly reflects mechanical effects of the surrounding cells.

We modeled the center cell as an *n*-sided regular frustum (*n* ≥ 3) with volume *V*_0_, apical surface area *A*_a_, and basal surface area *A*_b_. The set of *n*, *V*_0_, *A*_a_, and *A*_b_ uniquely determines the shape of the center cell. In this geometry, the center cell has an apical surface, a basal surface, or both and is adjacent to *n* first-neighbor cells with which it shares lateral boundary faces. The *n* first-neighbor cells also share lateral boundary faces and are aligned radially around the center cell. Additionally, we introduced several geometric parameters, including the height of the center cell *H*, apical perimeter of the center cell *P*_c_ (≥ 0), total area of the *n* boundary faces between the center cell and its first-neighbor cells *A*_c_ (≥ 0), and total area of the *n* boundary faces between the first-neighbor cells *A*_p_.

To describe the local deformation of the cell, we introduced the cell taper as defined by

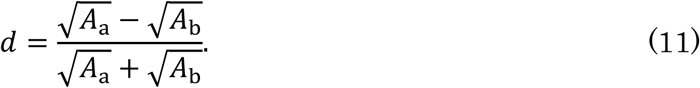

We modeled the mechanical energy of the epithelial sheet as follows. First, we assumed a fixed volume for each cell because the cell volume is almost constant during morphological changes in several organisms. Even under the constraints of constant cell volume and planar geometry, the cell-cell boundaries can move in principle, so *d* remained variable. To consider the same situation as a numerical simulation, we introduced the same energy function described in Equation 3.

The first term in Equation 3 is negligible because the total volume of each cell is assumed to be constant. However, the other terms are effective, because only the vertices constituting the center cell can move. We expanded the second term around the center cell and first-neighbor cells as *κ*(*A*_c_ + *A*_p_). Similarly, we expanded the third term as *Γ*{1 + (1/*n*)[1 − csc(*π*/*n*)]}, where {1 + (1/*n*)[1 − csc(*π*/*n*)]} is a correction factor that reflects the effects of the center cell and first-neighbor cells. Finally, we expanded the last term as *γ*[2 − csc(*π*/*n*)], where [2 − csc(*π*/*n*)] is a correction factor that reflects the effects of the center cell and first-neighbor cells. Thus, Equation 3 can be written as the mechanical energy of the center cell and first-neighbor cells in the planar sheet as described by

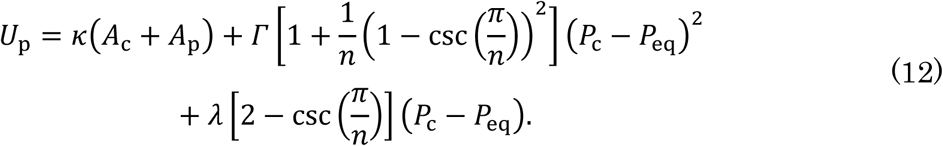

The packing geometry imposes geometric constraints on the physical parameters. We simultaneously solved Equation 13 and obtained *H*, *A*_a_, and *A*_b_ as functions of *ρ*, *V*_0_, and *d* as described by

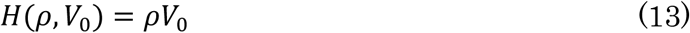

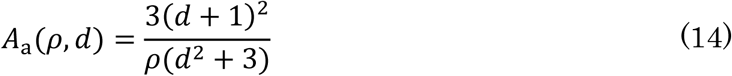

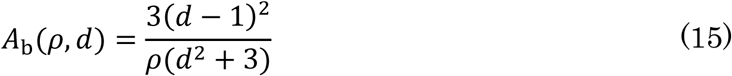

The geometric parameters *P*_c_, *A*_c_, and *A*_p_ in the energy function *U* are described as functions of *n*, *H*, *A*_a_, and *A*_b_ as described by

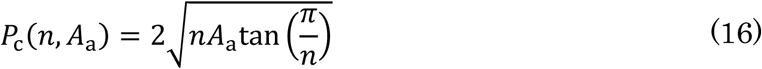

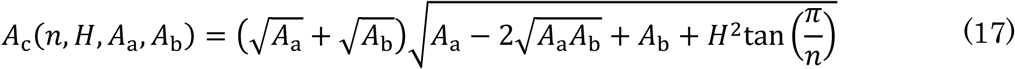

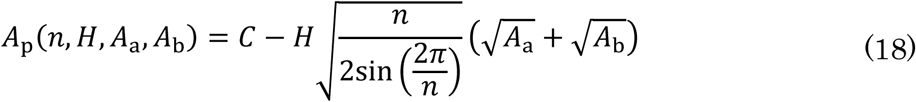

Where *C* is a constant that represents the total area of the boundary faces between neighbor cells and the cross section in the center cell. Because *A*_p_ is incorporated into the energy function in Equation 3 as a first-order process, *C* is negligible. Using the analytical expressions of *U*_p_, we then calculated energy landscapes as functions of *d* under specific *n*, *ρ*, and *λ* values. Physical parameters were nondimensionalized by units: length, *l*, was scaled by 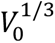, and energy, *ε*, by 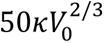.

### AFM measurement of cell stiffness

For the AFM measurements, we used a JPK Nanowizard 4 (Bruker) equipped as a confocal microscope (Expert line, Abberior). To measure stiffness at cell-cell junctions, we used a 3XC-GG cantilever (OPUS, spring constant 0.1–0.6 N m^-1^). Immediately before measurement, the medium was replaced with DMEM supplemented with 10% serum, 1% penicillin/streptomycin, and 10 mM HEPES. The following parameters were set: setpoint of 2–4 nN, *z* length of 6–8 µm, *z* speed of 100– 120 µm s^-1^, and a scan area of either 30 × 30 or 50 × 50 µm with a resolution of 128 × 128 or 256 × 256 pixels. After measurement, fluorescence images were captured using confocal microscopy, similar to the epithelial measurement. After averaging and baseline subtraction, Young’s modulus was calculated using a Hertz model with a triangular pyramid half-cone angle of 35°. The Young’s modulus map was then exported and converted to monochrome. Cell-cell junction regions were manually extracted from these maps, and the junctional stiffness at each pixel was obtained using Gwyddion software.

After each AFM measurement, 3D fluorescence images were acquired using a confocal microscope (Expert Line, Abberior) with a 60× lens (NA 0.7). Images were acquired with 560 nm illumination, 10% laser power, and a 10-µs exposure time per pixel. The scan size was 120 × 120 µm with a resolution of 600 × 600 pixels in the *xy* plane and 20–25 µm in the *z* axis with a 1 µm interval. To assess the dependence of epithelial stiffness, the planar cell density was calculated from the 3D fluorescence images. Because the AFM scan area was centered in the fluorescence image, the image was cropped to a size of 60 × 60 µm including the AFM scan area. A total of 19 stiffness maps were obtained from four dishes.

## Data Availability

Additional materials and data supporting the findings of this study are available from the corresponding author upon reasonable request.

## Code availability

Relevant code is available from the corresponding author upon reasonable request for academic use.

## Acknowledgements

We thank all the members of the Okuda Laboratory for discussions, A. Matsuoka for help with image analysis, and M. Nakayama, D. Wang, and H. Oshima at Kanazawa University for discussions. This work was supported by the WISE Program for Nano-Precision Medicine, Science, and Technology of Kanazawa University, Ministry of Education, Culture, Sports, Science and Technology (MEXT; to A.T.); the Japan Science and Technology Agency (JST), CREST [Grant No. JPMJCR1921, JPMJCR24B2]; the Japan Society for the Promotion of Science (JSPS), KAKENHI [Grant No. 21H01209, 21KK0134, 22H05170, 24H01398, 24H01937, 25K01118, 25K22468 (to S.O.), 24KJ0090 (to S.H.)]; the Japanese Agency for Medical Research and Development (AMED), the Program for Technological Innovation of Regenerative Medicine Grant [23bm0704065h0003 (to S.O.), 21bm0704060h0001, and 24bm1123049h0001 (to I.I.)], and the World Premier International Research Center Initiative, MEXT, Japan (to S.O.).

## Author Contributions

S.F. did data curation, formal analysis, investigation, methodology, software, validation, visualization, and writing (original draft and revised draft). R.D. did data curation, formal analysis, visualization. D. W., T.I., P.L., M.A., and A.T. did data curation, formal analysis, investigation, validation, and visualization. R.Y., K.O. did investigation, validation, supervision. S.H. formal analysis, investigation, methodology, software, validation. K.K. did resources and validation. A.S., I.I. did investigation. T.F., H.F. did funding acquisition, project administration, supervision. S.O. did conceptualization, data curation, funding acquisition, methodology, project administration, software, supervision, visualization, and writing (original draft and revised draft).

## Competing Interests

All authors declare no competing interests.

## Notes

### Competing Interest Statement

The authors have declared no competing interest.

### Summary of Updates

All text has been updated; Figures 4, 5, and 6 have been revised; and author affiliations have been updated.

